# Mouse trigeminal neurons respond to kokumi substances

**DOI:** 10.1101/540237

**Authors:** Sara C.M. Leijon, Nirupa Chaudhari, Stephen D. Roper

## Abstract

Using *in vi*vo confocal Ca^2+^ imaging, we investigated whether oral application of *kokumi* substances elicits responses in trigeminal somatosensory ganglion neurons of the mouse. Our results show that 100 μM γEVG (γ-Glu-Val-Gly), a potent *kokumi* stimulus, evokes responses in a very small fraction (0.6 %) of neurons in area V3 (oral sensory field) of the trigeminal ganglion. By comparison, cooled artificial saliva elicited thermal-evoked responses in >7 % of V3 trigeminal ganglion neurons. γEVG-evoked responses were small and quite variable, with latencies ranging from 2 to over 200 sec. Co-application of the calcium sensing receptor (CaSR) inhibitor NPS-2143 significantly decreased γEVG-evoked activity. Furthermore, we show that 4 additional *kokumi* substances evoked responses in mouse trigeminal ganglion neurons. All neurons responding to *kokumi* compounds were small cells, with mean diameters below 20 μm. In summary, our data show that certain physiological and pharmacological properties of responses to *kokumi* compounds can be recorded from sensory neurons in the trigeminal ganglion of living mice. Thus, sensory neurons in the somatosensory trigeminal ganglia may transmit signals from the oral cavity to the central nervous system to generate the texture perceptions that are part of the enigmatic sensations evoked by *kokumi* substances.

## 2. Introduction

Sensations elicited by *kokumi* substances are difficult to define and to quantify. It is often described as “mouthfulness”, “thickness”, and “continuity”—characteristics that cannot readily be explained in terms of taste alone. *Kokumi* compounds, though themselves evoking no taste, are also claimed to enhance sweet, salty, and umami tastes (Ohsu et al, 2010). These features are not easily measured and often rely on human sensory panels and verbal descriptions. Finding an animal model that can be used to test and quantify *kokumi* sensations would be valuable for research and development of *kokumi* compounds. This chapter outlines progress along those lines and describes a new approach to analyzing koku taste in a mouse model.

Afferent somatosensory neurons in the trigeminal ganglion innervate the tongue and oral surfaces. These neurons are sensitive to many different modalities, including mechanical, thermal, chemesthetic, and nociceptive stimuli. Trigeminal sensory neurons may thus participate in transmitting sensations such as texture and viscosity, which could be perceived as mouthfulness, thickness, and continuity. Interestingly, trigeminal sensory neurons in the rat are reported to express a calcium sensing-receptor, CaSR (Heyeraas et al, 2008), which has been shown to be activated by *kokumi* substances (Oshu et al, 2010; Amino et al, 2016). Collectively, these features provide a rationale for studying the involvement of trigeminal ganglion somatosensory neurons in responses to *kokumi* compounds. Accordingly, we have taken advantage of a powerful new experimental approach that combines Ca^2+^ imaging, genetically-engineered mice, and *in vivo* scanning laser confocal microscopy (Fig. 1) to investigate responses of mouse trigeminal ganglion neurons to oral stimulation with *kokumi* stimuli. Our aim was to determine whether trigeminal ganglion neurons respond to *kokumi* substances that are presented in the oral cavity, and in particular, to stimulation by the prototypic koku substance, γ-glutamyl-valinyl-glycine (γEVG). Using this animal model, we tested sensory neuron responses to γEVG in the living animal and characterized some of the properties of this *kokumi* compound. Our particular interests were in the viscous sensations elicited by γEVG and in the identity of possible membrane receptors for that compound.

**Figure 1.**
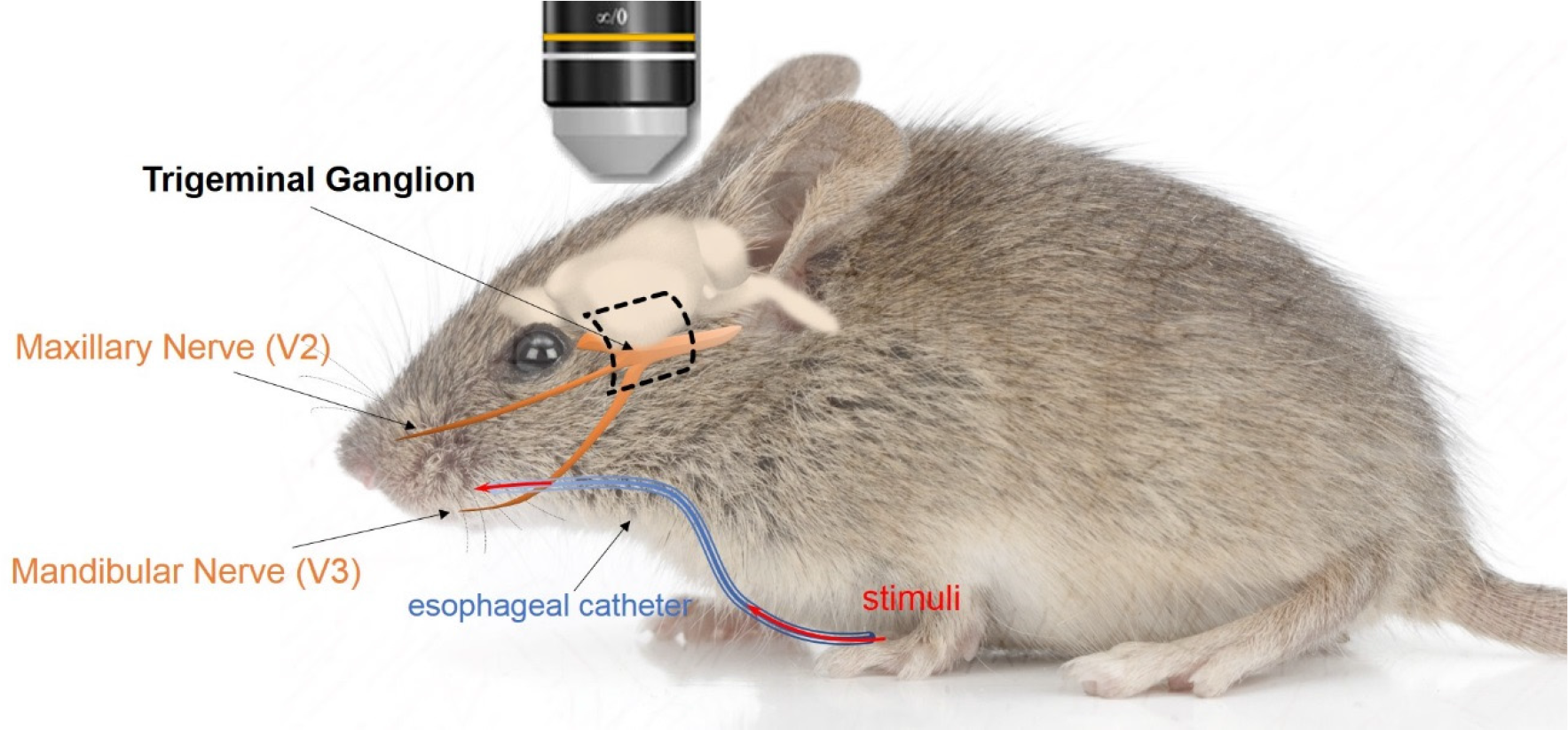
Schematic of the trigeminal ganglion location in the mouse. The trigeminal ganglion (orange) resides at the bottom of the cranium, below the brain. The mandibular and maxillary branches of the trigeminal nerve innervate the oral cavity. A surgery is performed to carefully expose the trigeminal ganglion via a dorsolateral window (dashed rectangle). The surgically prepared mouse is thereafter transferred to a confocal laser scanning microscope, and neuronal calcium responses are recorded with a 10x long-working distance objective.

## 3. Methodology

### 3.1 Animal model

We used transgenic mice that express the calcium indicator GCaMP3 or GCaMP6s in sensory neurons, including those in the trigeminal ganglion (Leijon et al, 2019). GCaMP3 mice were obtained from X. Dong, Johns Hopkins and express GCaMP3 as a knock-in/knockout at the Pirt locus (Kim, Chu et al, 2014). We also used floxed GCaMP6s (Jax #024106) crossed with Pirt-Cre mice (X Dong, Johns Hopkins) to generate mice in which sensory neurons expressed GCaMP6s. All mice were backcrossed to C57Bl/6 for 8-10 generations and adult mice of both sexes were used. Animals were housed with a 12-hour light cycle and food and water *ad libitum* and all experiments were carried out during the daylight cycle. All procedures for surgery and euthanasia were reviewed and approved by the University of Miami IACUC.

### 3.2 Surgical preparation

Mice were anesthetized with ketamine and xylazine (intraperitonially 120 mg/kg ketamine, 10 mg/kg xylazine) and anesthetic depth was monitored by the hind paw withdrawal reflex. Ketamine booster injections were administered to ensure continued surgical plane of anesthesia throughout the surgery and imaging session. The animal’s core temperature was continuously monitored with a rectal probe and maintained between 35 and 36 °C.

For the initial stage of surgery, the anesthetized mouse was placed supine on a far infrared surgical warming pad (DCT-15, Kent Scientific). The trachea was exposed and cannulated for facilitated respiration during oral stimulus presentation. A flexible tube was inserted through the esophagus to produce a uniform stimulus delivery into the oral cavity (Sollars and Hill 2005; Wu, Dvoryanchikov et al. 2015; Leijon et al, 2019). Next, the mouse was placed in prone position for surgical exposure of the region of the trigeminal ganglion that innervates the oral cavity, V3 (Fig. 2). Muscle tissue on the dorsolateral side of the skull was cauterized and the zygomatic arch removed. A small cranial window was drilled open and hemispherectomy was performed by careful aspiration to allow optical access to the dorsal part of the trigeminal ganglion. The head was stabilized by affixing a custom head holder to the skull with a nylon screw and dental acrylic. To maintain a favorable neuronal environment, from the moment of exposure the ganglion was intermittently rinsed with Tyrode’s solution at 35 °C.

**Figure 2.**
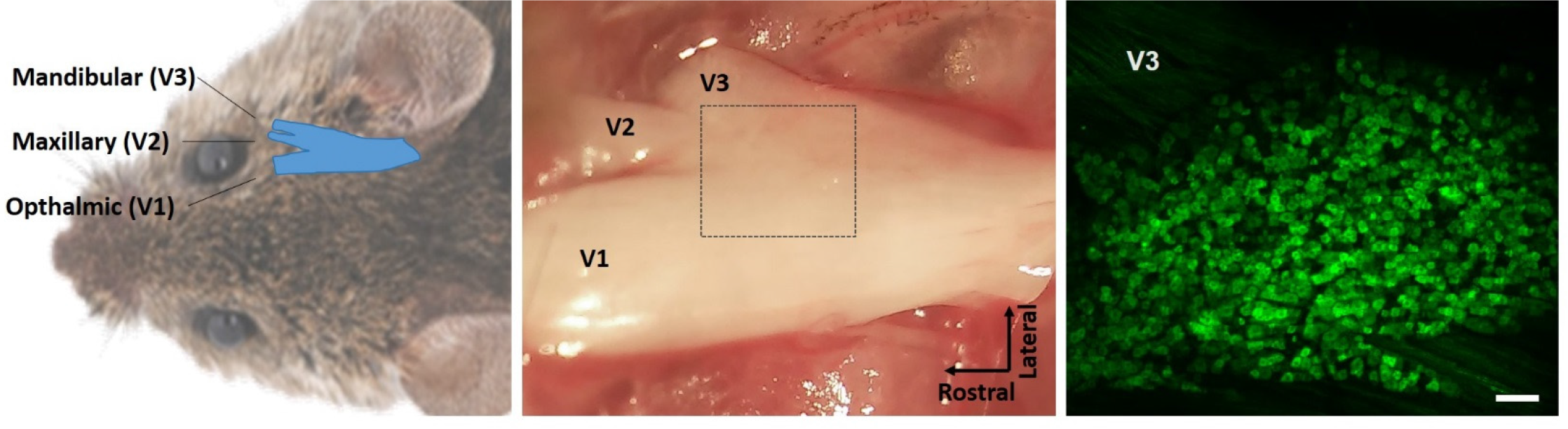
Location and surgical view of the trigeminal ganglion. a) Dorsal view, displaying the relative position of the trigeminal ganglion (blue). b) photograph showing V3 region of the surgically exposed trigeminal ganglion and the area where functional imaging was carried out (rectangle). c) Under the confocal laser scanning microscope, the calcium indicator, GCaMP, in trigeminal ganglion neurons fluoresces green. Stimulus-evoked responses generate robust fluorescence increases in stimulated neurons. A typical recording field contains around 1000 ganglion neurons and can be imaged for over 5 hours. Scale bar, 100 μm.

### 3.3 Functional *in vivo* confocal calcium imaging

The surgically-prepared mouse was transferred to the stage of an Olympus FV1000 confocal microscope equipped with a 10x-long working distance objective, (UPlanFl, N.A. 0.3). Confocal scans of GCaMP3-/GCAMP6s-labeled ganglion neurons were taken at ∼1 hz using 488-nm laser excitation with a 505–605 nm emitter filter. Stimuli were applied as 1 mL over 10 seconds, delivered retrogradely via the esophageal tube. When temperature stimuli were applied, the resulting oral temperature was measured with a thermal probe connected to a wireless transmitter (UWBT-TC-UST-NA, OMEGA Engineering, Inc.).

### 3.4 Reagents

All reagents were purchased from Sigma apart from *kokumi* substances, which were obtained from Ajinomoto Co. Inc. These included glutathione (GSH; CAS 70-18-8), γ-glutamyl-alpha-aminobutyrate (γ-Glu-Abu; CAS 16869-42-4), γ-glutamyl-valinyl-glycine (γ EVG, γ-Glu-Val-Gly; CAS 38837-70-6), poly-L-lysine (CAS 28211-04-3), cinacalcet-HCl (CAS 364782-34-3), and the Calcium-Sensing Receptor inhibitor NPS-2143. Stock solutions of *kokumi* substances were dissolved in H_2_0 and stored at 4° C. On the experimental day, *kokumi* stimuli were diluted to working solutions in artificial saliva containing 15 mM NaCl, 22 mM KCl, 3 mM CaCl2, 0.6 mM MgCl2, pH 5.8.

### 3.5 Data analysis

Optical scans of neuronal fluorescence were digitized using Fluoview software (Olympus) and videoimages were stabilized using FIJI (ImageJ). Baseline-subtracted videos were used to identify areas with potential neuronal responses and regions of interest (ROIs) were manually drawn over individual ganglion neurons. ROIs were analyzed with MatLab (Mathworks) using custom code, modified from Wu et al., 2015, to correct for any small baseline drifts. Calcium transients were quantified as peak stimulus-evoked fluorescence change (Δ*F*) divided by baseline fluorescence (*F*_0_), i.e. ΔF/F_0_. The criterion for identifying responses was ΔF/F_0_ > 5 standard deviations above baseline (Fig 3). For thermal stimuli, criteria also included that responses occurred at consistent latencies after stimulus onset. A single field of view, within the V3 area of the trigeminal ganglion, was analyzed for each experimental animal. At the end of an experiment, the animal was euthanized by CO_2_ followed by cervical dislocation.

**Figure 3.**
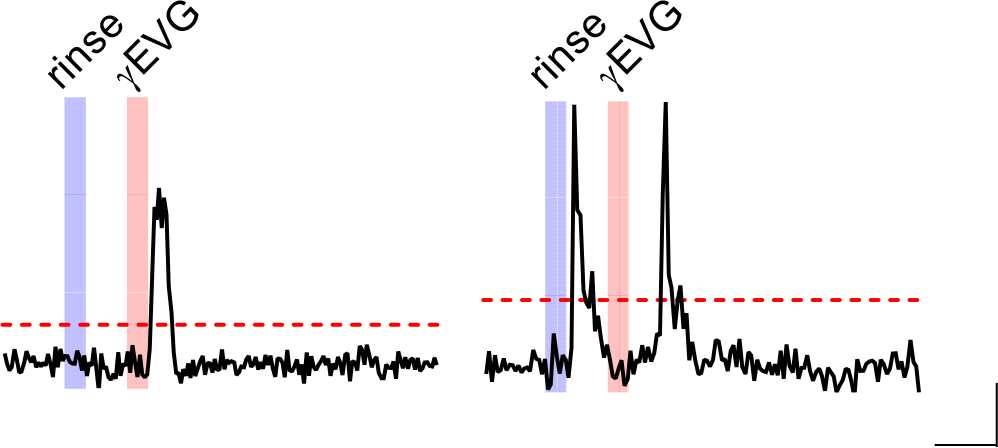
Examples of responses (ΔF/F_0_) from two separate neurons to oral stimulation with 100 μM γEVG. Here, as in later figures, stimuli were superfused into the oral cavity for 10 seconds (shaded areas). Red dashed lines mark criterion level above which ΔF/F_0_ must rise to be designated as responses (i.e., ΔF/F_0_ > 5 times the standard deviation above baseline, see Methods). Ca^2+^ transients are seen after γEVG stimulation (orange shaded area) in both neurons. However, only the neuron on the left is confirmed to be γEVG-responsive. The neuron on the right responded equally to the control stimulus (artificial saliva, blue shaded area) as to γEVG, likely reflecting responses to temperature or flow (which always appear as single, transient responses with brief latencies, see Fig. 7 and Leijon et al, 2019). Calibs, 30 sec, 0.5ΔF/F_0_.

Unless otherwise specified, statistical significance was determined using two-tailed Student’s t-test between two groups, otherwise by unpaired or paired one-way ANOVA, using Prism v.6 (GraphPad). Statistical significance was defined as p<0.05.

## 4. Results

### 4.1 *Kokumi* peptide γ-Glu-Val-Gly

#### 4.1.1 Trigeminal primary afferent neurons respond to the *kokumi* peptide γ-Glu-Val-Gly

To investigate if oral exposure to γ-Glu-Val-Gly (γEVG) elicits trigeminal responses *in vivo*, the oral cavity of the mouse was superfused with 100 μM γEVG. Because many trigeminal ganglion neurons are also sensitive to other sensory stimuli, for example temperature (Yarmolinsky et al, 2016; Leijon et al, 2019), γEVG was preceded with a control stimulus consisting of artificial saliva at the same temperature and perfusion rate as the γEVG application (Fig 3). Our results show that γEVG (100 μM) applied orally elicits responses in a small population of trigeminal ganglion neurons (Fig. 4).

**Figure 4.**
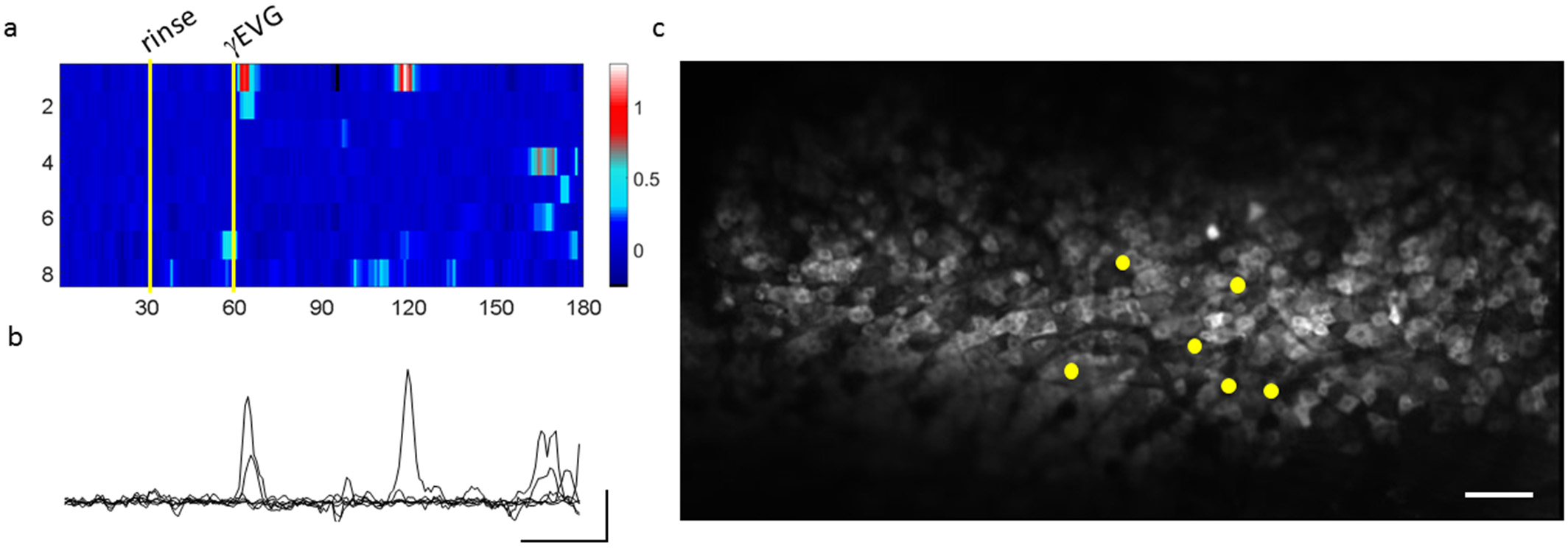
Oral γEVG evokes responses in trigeminal ganglion neurons. a) A heat map displaying the fluorescence change (ΔF/F_0_; color bar) of 8 trigeminal ganglion neurons over a 180 second recording. Yellow lines indicate onset of stimuli (stimulus duration = 10 sec). The top 6 neurons show responses to 100 μM γEVG. Note that these neurons are silent during the baseline (first 30 seconds), and are unaffected by the control (artificial saliva) stimulus. The bottom 2 cells show nonspecific activity and are not considered γEVG-responsive. b) Superimposed traces of the 6 γEVG-responsive neurons from recording shown in a. Calibs, 30 sec, 0.5 ΔF/F_0_. c) Location of the 6 γEVG-responsive neurons in the trigeminal ganglion. Note that imaging the ganglion allows simultaneous recording of a large number of trigeminal ganglion neurons (typically around 1000). Scale bar, 100 μm.

#### 4.1.2 Characteristics of responses evoked by γ-Glu-Val-Gly

γEVG responses in trigeminal neurons had variable latencies. Although occasionally we observed well-defined, transient γEVG-evoked responses within a few sec of the stimulus onset (Fig 5a), more often γEVG responses had much longer latencies and were characterized by prolonged bursts of activity (Figs 4, 5b-c). Across a sample of 5 mice (26 cells), the mean latency of γEVG responses was 78±12 seconds (Fig 6a). The average amplitude was 1.4±0.2 ΔF/F_0_ (Fig 6b).

**Figure 5.**
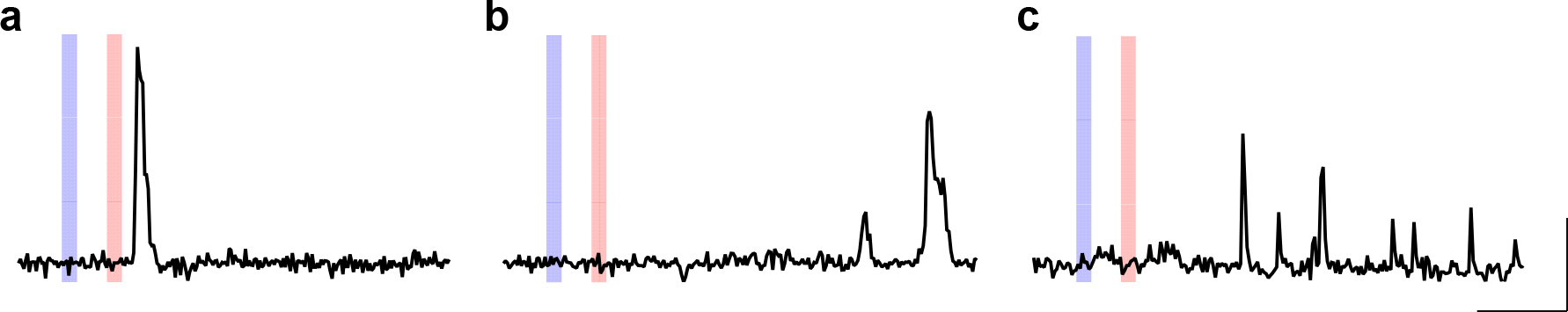
Oral γEVG evokes short- and long-latency responses in trigeminal ganglion neurons. Traces show evoked responses in 3 different neurons to 100 μM γEVG applied orally for 10 sec (orange shaded areas). Responses consisted of a single transient (a), or multiple transients (b,c). Note that an initial rinse with artificial saliva for 10 sec (blue shaded areas) did not elicit responses in any of these neurons. Calibs, 60 sec, 1 ΔF/F_0_.

**Fig 6.**
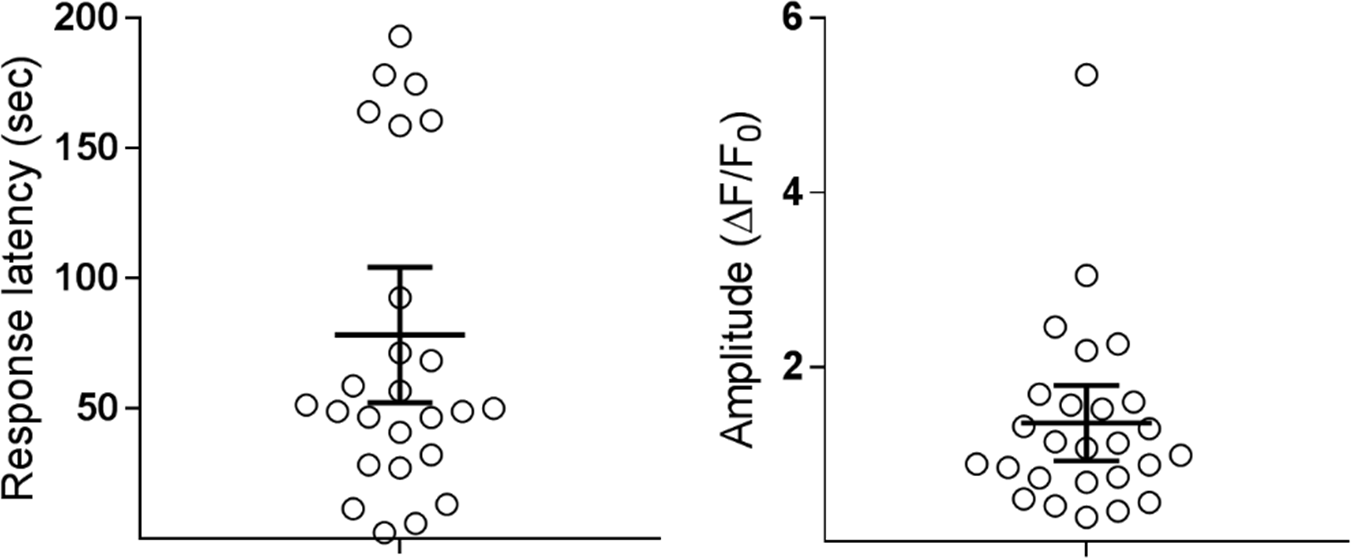
Properties of γEVG-evoked responses. The response latencies (sec) and amplitudes (max ΔF/F_0_) were measured across a sample of 5 mice (26 cells). Error bars show mean ± 95 % CI.

#### 4.1.3 Comparing γ-Glu-Val-Gly responses to thermal responses

The incidence of trigeminal ganglion neurons stimulated by oral application of γEVG was quite low (0.6±0.0 %, 3 mice). By comparison, the incidence of trigeminal ganglion neurons that responded to orostimulation with cooled artificial saliva (∼15 °C) was > 10 fold higher, 7.4±0.6 %, a significant difference (3 mice; p<0.0001; Fig 7b). From the same sample, 36 % (8/22) of γEVG-responsive neurons were also sensitive to cooling the oral cavity.

**Figure 7.**
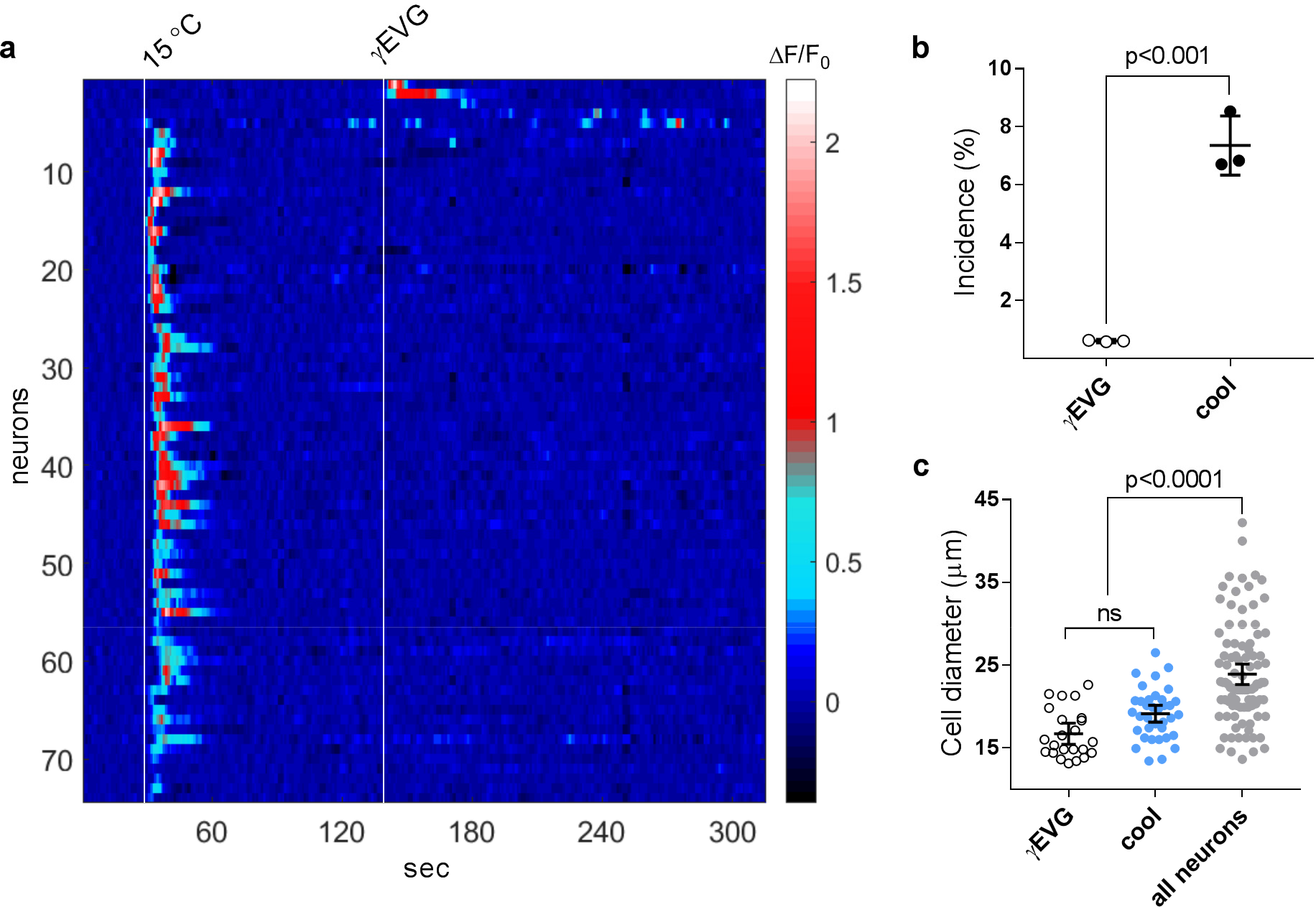
Comparing γEVG responses to responses evoked by cooling the oral cavity. a) Heat map showing the fluorescence change (ΔF/F_0_; color bar) of 76 trigeminal ganglion neurons during 10 sec oral stimulation (onset at white lines) with cold (∼15 °C) artificial saliva and with 100 μM γEVG, sequentially. b) incidences for γEVG-responding and cool-sensitive neurons (3 mice, 22 and 272 neurons, respectively; Students t-test). c) Cell sizes were measured for γEVG-responsive (7 mice, 23 neurons) and cool-sensitive neurons (3 mice, 36 neurons), as well as for a random sampling of all trigeminal neurons (3 mice, 94 neurons). Error bars show mean ± 95 % CI.

The trigeminal ganglion consists of a diverse population of sensory neurons (e.g. mechanosensitive, thermosensitive, nociceptive, itch) with varying cell sizes. The mean cell diameter (Fig 7c) of γEVG-responding neurons was 16.7±0.61 μm, which is significantly smaller than the average of all trigeminal neurons (23.9±0.63, p<0.0001). Cool-sensing neurons, as well, had small diameters (19.1±0.51 μm, p=0.19 μm).

#### 4.1.4 Effect of CaSR-blocker NPS 2143 on γ-Glu-Val-Gly-evoked *kokumi* responses

CaSR is thought to be involved in responses evoked by *kokumi* substances observed in HEK-293 cells transiently expressing human CaSR (Oshu et al, 2010), in CaSR-expressing taste cells in lingual slices (Maruyama et al, 2012), and in human sensory analyses (Kuroda & Miyamura, 2015). Thus, we investigated whether the CaSR inhibitor NPS-2143 had an effect on γEVG-evoked responses recorded *in vivo* from trigeminal ganglion neurons. We recorded responses to oral stimulation with γEVG alone or in combination with 30 μM NPS 2143 (Fig. 8). Because γEVG often elicited multiple calcium transients (Fig. 5), we calculated the area under curve (AUC) to quantify γEVG responses. The results showed that NPS significantly decreased γEVG responses (p<0.0001) (Fig. 8). NPS had no effect on stimulation with artificial saliva (p>0.05, Fig. 8a). To exclude the possibility that the decline in γEVG responses following NPS treatment was merely due to “rundown” during prolonged recordings, we conducted a second series of experiments where γEVG alone was applied twice in the absence of the CaSR inhibitor (Fig. 8b). Importantly, there was no decrease in AUC of γEVG responses (p=0.29). Furthermore, there was no difference between AUC measurements from γEVG alone responses between the first and second of these series of experiments (p=0.5589). These data suggest that γEVG-evoked responses in the trigeminal ganglion neurons can be inhibited by applying the CaSR inhibitor NPS 2143 into the oral cavity.

**Figure 8.**
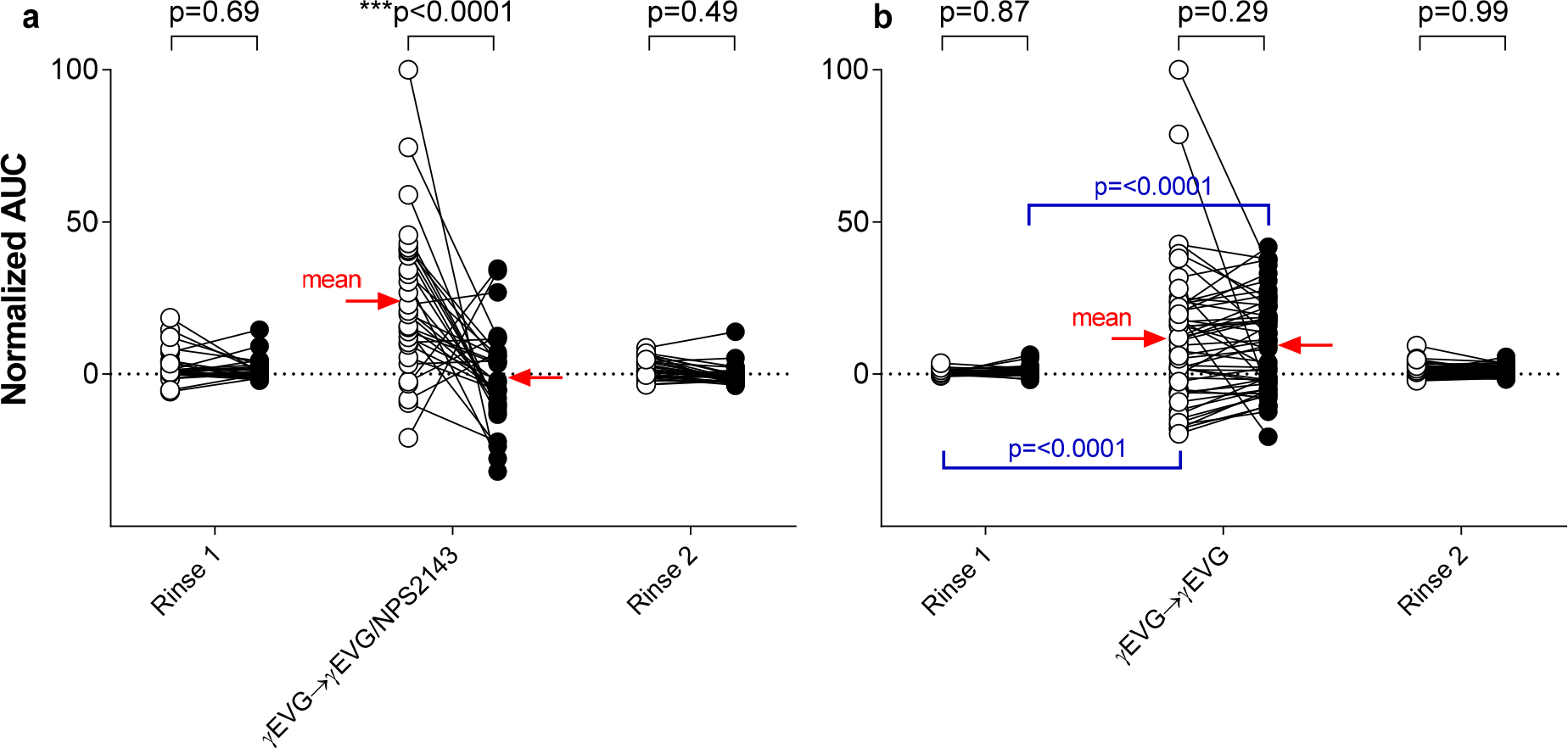
The CaSR inhibitor NPS 2143 reduces γEVG responses in trigeminal ganglion neurons. a) A sequential series of oral stimuli: artificial saliva (rinse 1) → 100 μM γEVG → artificial saliva (rinse 2) was applied and trigeminal neuron responses (area under the curve of ΔF/F_0_, AUC) were recorded. After this control series was completed (open symbols), the oral cavity was flushed with 30 μM NPS 2143 and the same sequence of stimuli was repeated (closed symbols). The response to γEVG was significantly depressed in the presence of NPS 2143 (closed symbols; paired t-test). Responses to artificial saliva, although small, were unaffected by NPS 2143 (2 mice, 31 neurons). b) In a second series of experiments, the same protocol was repeated but without NPS 2143 to test for rundown of γEVG responses during the prolonged recording and repeated stimulations. No rundown of γEVG responses occurred (3 mice, 51 neurons).

### 4.2 Effect of viscosity on γ-Glu-Val-Gly-evoked responses

#### 4.2.1 γ-Glu-Val-Gly & Xanthan gum

In human sensory analyses, *kokumi* substances stimulate a sensation of enhanced thickness and mouthfeel during food intake (Oshu et al, 2010). We investigated whether a viscous solution evoked neuronal responses and whether viscosity affected γEVG responses. Xanthan gum is a common commercial food thickener. Xanthan gum alone (0.5%) appeared to evoke responses in trigeminal ganglion neurons but the effects were diffuse and difficult to quantify (Fig 9). To analyze the data we removed data (ΔF/F_0_) that was did not reach a level of 5 times the standard deviation above baseline and calculated the area under the remaining signal (AUC). We also counted the number of peaks in ΔF/F_0_ (Fig 9c). The results show that oral xanthan gum alone evoked increased activity in trigeminal ganglion neurons. Stimulating with a combination of γEVG and xanthan gum resulted in a somewhat larger AUC for the combination response than that evoked by γEVG alone, but the difference was not statistically significant (p=0.51). The same result was obtained for the number of peaks (ΔF/F_0_) (p=0.56). Thus, oral application of the food thickener xanthan gum can evoke responses in the trigeminal ganglion, but an interaction between xanthan gum and the *kokumi* substance γEVG cannot be confirmed.

**Figure 9.**
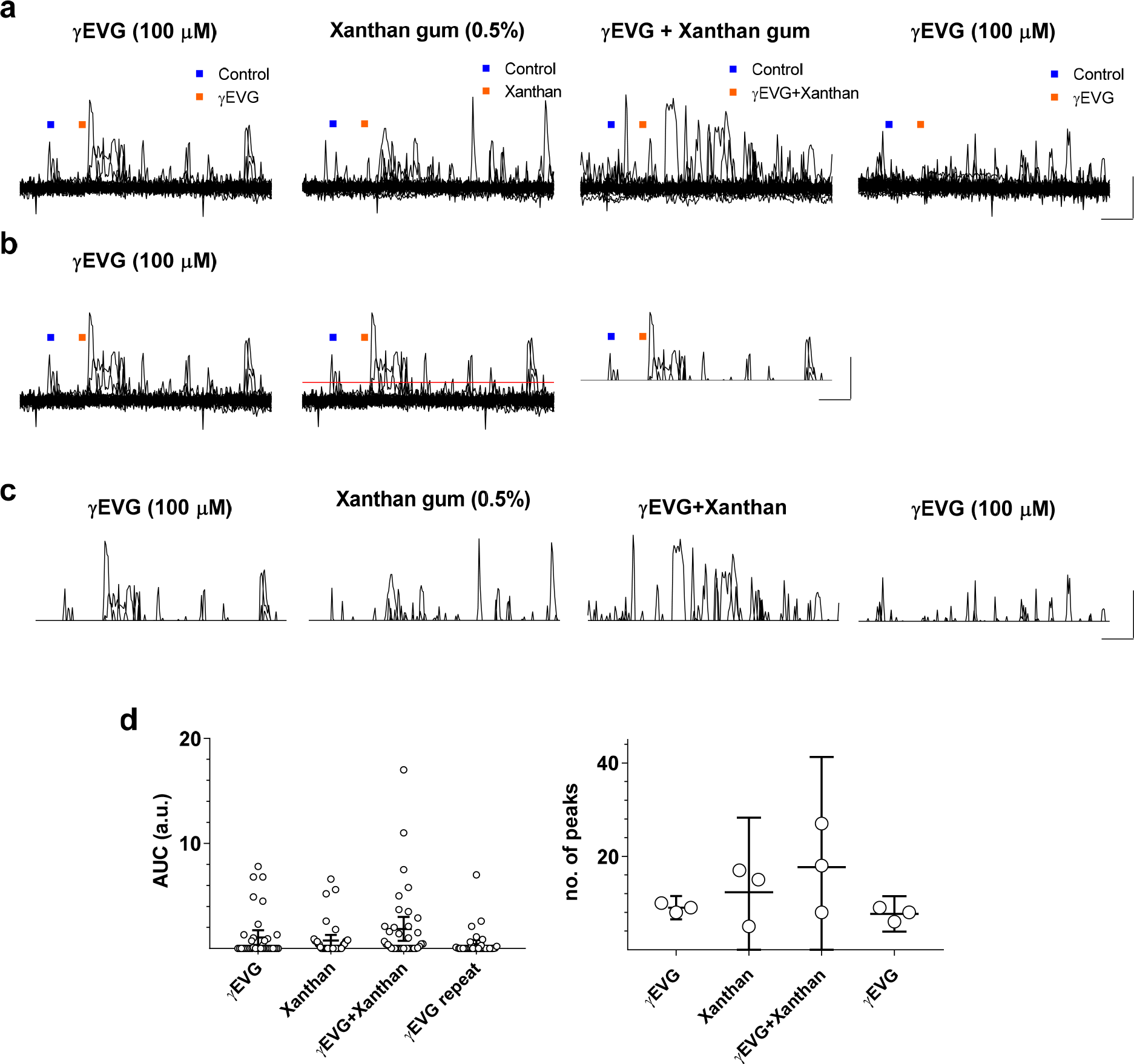
The effect of the food thickener Xanthan gum on γEVG-evoked responses. a) Original recordings during oral application of γEVG and Xanthan gum individually, as well as combined. b) As exemplified here for a response to γEVG (left trace), for each recording we calculated the signal criterion (equal to > 5 s.d. above baseline, red line in middle trace) and eliminated any points less than this value (right trace). c) The responses that reached the cutoff criterion were analyzed for area under the curve (AUC) as well as number of peak values of ΔF/F_0_. d) Left, each data point corresponds to AUC for responses noted on the x axis (38 neurons, 3 mice). Right, each data point corresponds to the total number of peaks in ΔF/F_0_ traces, per mouse (3 mice). Error bars show mean + 95 % CI. Calibs, 30 sec, 1 ΔF/F_0_.

#### 4.2.2 γ-Glu-Val-Gly & glucomannan

Next, we tested if another thickener, glucomannan, had an effect on γEVG-evoked responses. Similar to Xanthan gum, oral stimulation with glucomannan alone (0.3%) evoked responses in trigeminal ganglion neurons (Fig 10a). There was no interaction between γEVG and glucomannan. Stimulating the oral cavity with a combination of γEVG and glucomannan did not evoke responses that differed from stimulating with γEVG alone (no difference in AUC of the responses, p=0.37, or in number of peaks, p=0.65) (Fig 10b).

**Figure 10.**
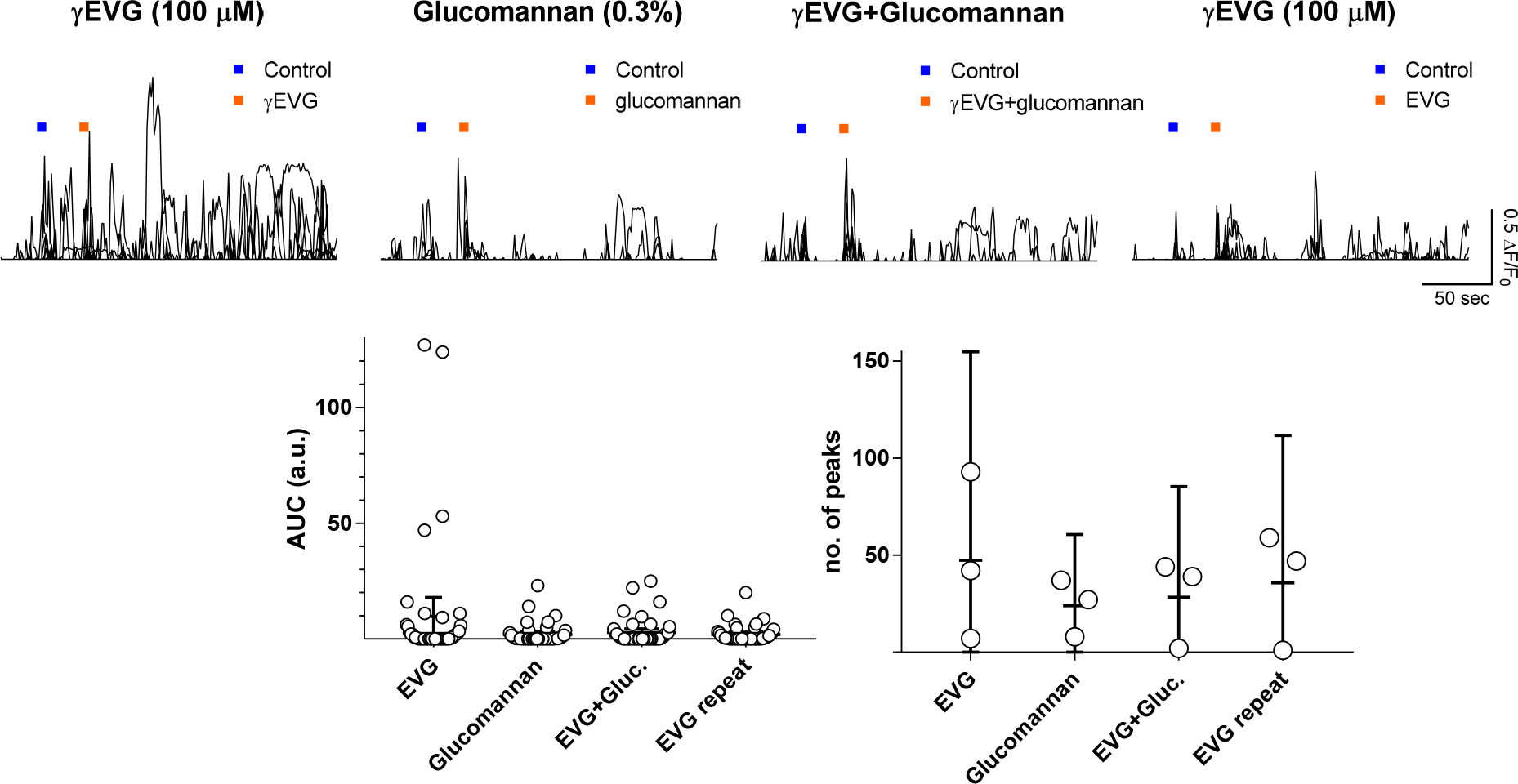
The effect of glucomannan on γEVG-evoked responses. a) As for Xanthan gum in Fig 9, responses above the cutoff criterion were analyzed for AUC as well as number of peaks in the trace. b) Left, each data point corresponds to AUC (45 neurons, 3 mice). Right, each data point corresponds to the total number of peaks in the responses, per mouse (3 mice). Responses to glucomannan and γEVG alone and in combination did not differ in either AUC or number of peak responses. Error bars are mean + 95 % CI.

#### 4.2.3 Sizes of: γEVG-responding neurons versus neurons responding to thickeners

In the thickener experiments, some neurons showed responses to both γEVG and thickener (xanthan gum/glucomannan). The average diameter of γEVG-responding neurons was 18.2±0.7 μm (Fig. 11), which is not signficantly different from our previous measurements (Fig. 7 p=0.13). The average sizes of neurons responding to xanthan gum (17.2±1.6 μm) or glucomannan (16.1±0.5 μm) were similar.

**Figure 11.**
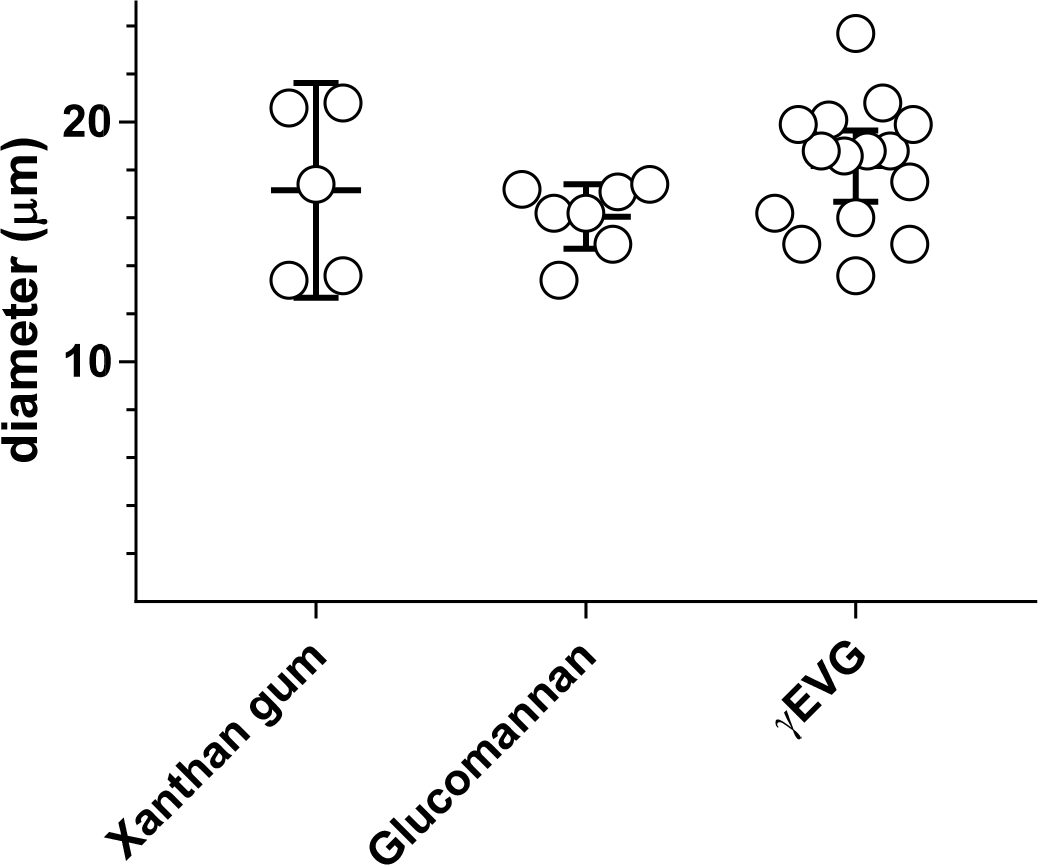
The cell diameter (μm) of neurons that respond to viscous stimuli or to γEVG. There was no difference in the diameters between trigeminal ganglion neurons responding to oral application of xanthan gum (3 mice, 5 cells), glucomannan (3 mice, 7 neurons) or γEVG (3 mice, 17 neurons).

### 4.3 Additional *kokumi* substances

Although γEVG has been described as a potent *kokumi* substance (Kuroda and Miyamura, 2015), there are other peptides with *kokumi*-like activity (Amino et al, 2016)(Kuroda and Miyamura, 2015). We investigated if *kokumi* substances other than γEVG elicited Ca^2+^ responses in mouse trigeminal ganglion neurons. Specifically, we tested glutathione (GSH/γ-Glu-Cys-Gly; 4 mice), γ-Glu-Abu (2 mice), poly-L-lysine (2 mice), CaCl_2_ (2 mice), and the calcium-mimetic drug Cinacalcet (3 mice).

#### Glutathione (GSH)

The effect of GSH on taste has been known since the early 1990’s when it was first extracted from garlic water and shown to enhance umami intensity (Ueda et al, 1990). This “intensifying” effect was termed *koku*. GSH has since been shown to also affect other taste qualities (e.g., Oshu et al, 2010). We found that oral application of 1 mM GSH evoked responses in mouse trigeminal ganglion neurons. The responses (Fig 12a) were of similar latency and duration as those of γEVG (Fig 6). Across 4 mice, GSH evoked responses in 26 trigeminal ganglion neurons.

**Figure 12.**
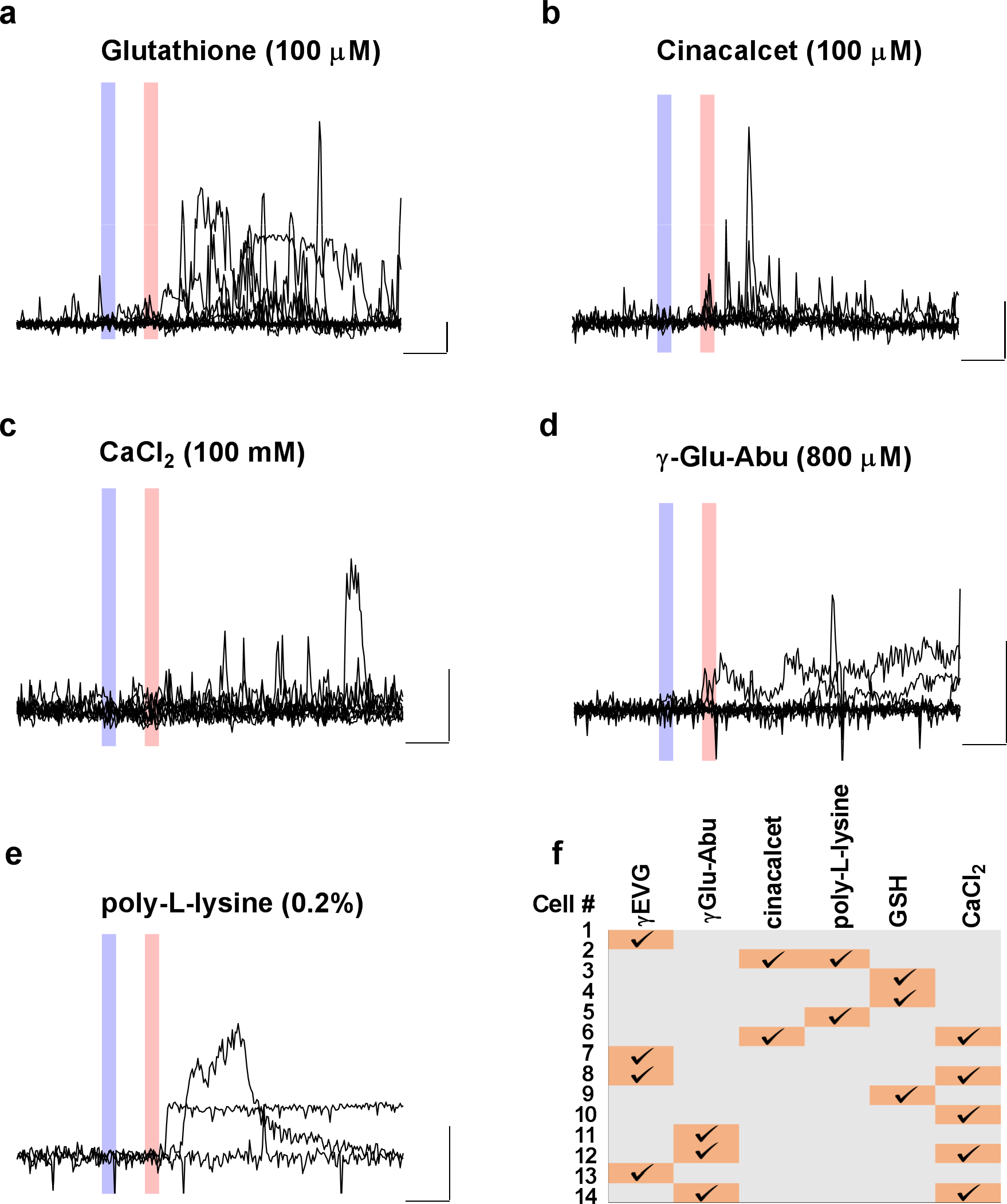
*In vivo* trigeminal ganglion neuron responses elicited by 5 selected *kokumi* substances applied orally. In a-e, each graph contains data from one mouse for which all responding neurons to the respective substance have been overlaid. Shaded areas correspond to the stimuli onsets of artificial saliva (blue) and the respective *kokumi* substance (orange). Note that the control stimulus (artificial saliva) did not elicit responses. f) One experiment was conducted where all 6 *kokumi* substances were applied sequentially. A total of 14 neurons (out of >1000) responded to at least one *kokumi* substance, and 5 of those 14 neurons responded to more than one *kokumi* substance. Calibs, 30 sec, 1 ΔF/F_0_.

#### Cinacalcet

Cinacalcet acts as a calcimimetic and causes allosteric activation of CaSR. In human sensory analyses, oral 38 μM cinacalcet showed significant *kokumi*-like taste enhancement (Oshu et al., 2010). We found that 100 μM cinacalcet evoked responses in the trigeminal ganglion (Fig 12b). Across 3 mice, cinacalcet evoked responses in 12 trigeminal ganglion neurons.

#### CaCl_2_

A certain basal level of calcium is needed for *kokumi* substances to activate CaSR (Wang et al., 2006; Oshu et al., 2010), and calcium salts have *kokumi*-like activity (Oshu et al., 2010). We superfused the oral cavity with 100 mM CaCl_2_ and observed trigeminal neuron responses that were similar in delay and characteristics to those of γEVG (Fig 12c). Across 2 mice, CaCl_2_ evoked responses in 11 neurons.

#### γ-Glu-Abu

The *kokumi* substance, γ-glutamyl-α-aminobutyrate (γ-Glu-Abu) is a natural substrate of glutathione synthetase, also known under its systematic name butanoic acid. It one of the most potent dipeptide *kokumi* substances (Amino et al., 2016). (Fig 4.9d). Across 2 mice, γ-Glu-Abu (800 μM) evoked responses in 11 neurons (Fig 12d).

#### Poly-L-lysine

Polylysine is a basic peptide that activates CaSR (Brown & MacLeod, 2001). In human sensory analyses, 0.08 % polylysine has been shown sufficient to enhance taste (Oshu et al, 2010). We found that, in the mouse, oral stimulation with polylysine (0.2 %) elicited responses only in a few trigeminal neurons. Polylysine responses were quite prolonged (Fig 12e). Across 2 mice, a total of 6 neurons were responsive to polylysine at this concentration.

### 4.4 Overlap between *kokumi* substances

To determine if the above *kokumi* substances elicited responses in identical, overlapping, or separate groups of trigeminal ganglion neurons, we applied each compound alone and sequentially in one experiment. The results indicated that 9 of the 14 neurons responded to one compound only; 5 neurons responded to 2 compounds (Fig 12f). There was no obvious pattern in the overlap. No consistent population of “*kokumi* sensing” trigeminal ganglion neurons was evident.

## 5. Discussion

In this study, we investigated responses of mouse trigeminal ganglion neurons to oral stimulation with *kokumi* compounds. We used transgenic mice that express GCaMP in sensory neurons to conduct *in vivo* imaging and analyze a number of *kokumi* compounds as well as testing the effect of blocking CaSR, a putative receptor for *kokumi* compounds.

Trigeminal sensory neuron responses to γEVG (γ-Glu-Val-Gly) had variable latencies (2 to >200 seconds) and were often characterized by prolonged activity. This likely reflects the gradual penetration of the compound into the lingual epithelium where the trigeminal afferent sensory nerve endings are situated. These trigeminal terminals can be quite superficial in the lingual epithelium (Leijon et al, 2016). During sensory evaluations, panelists report that γEVG added to food enhances taste intensity, peaking at 10 seconds but still present up to 40 seconds (Oshu et al., 2010). Those data would be consistent with the present study where oral stimulation with γEVG evoked long-lasting responses in trigeminal ganglion neurons.

CaSR has previously been implicated as a mechanism of action for *kokumi* taste both *in vitro* (Amino et al, 2016) and in human sensory analyses (Oshu et al, 2010). Thus, we tested if the CaSR inhibitor NPS-2143 affected γEVG responses. Indeed, co-application of NPS-2143 significantly decreased γEVG-evoked activity. Whether NPS-2143 selectively and specifically depresses only γEVG responses and not other stimulus-evoked activity (e.g. mechanical, thermal, or nociceptive) remains to be tested. Also, we did not test NPS-2143 on responses to other *kokumi* compounds such as glutathione. Moreover, it is unclear to what extent CaSR is expressed in the trigeminal ganglion. RNAseq analyses revealed CaSR expression only in few (<1%) of mouse trigeminal ganglion neurons, and only at negligible levels (Nguyen et al, 2017). Yet if CaSR is only expressed in neurons that innervate the oral mucosa, this might explain the low percentage trigeminal ganglion neurons that express this receptor. It would not, however, explain the low level of CaSR expression per cell that Nguyen et al (2017) report. Taken collectively, our findings do not offer conclusive evidence for the involvement of CaSR in trigeminal neurons responses to *kokumi* substances.

Based on the *koku* characteristics “mouthfullness” or “thick flavor”, we studied whether there is a sensory interaction between *kokumi* compounds and food thickeners. The data did not reveal significant interactions between oral γEVG and xanthan gum or glucomannan. Because a population of trigeminal sensory ganglion neurons responded to oral stimulation with γEVG or to xanthan gum (or glucomannan) alone, the perception of *koku*-enhanced thickness that sensory panels report (Oshu et al, 2010) may be generated by *kokumi* compounds stimulating viscosity-sensitive trigeminal sensory afferent fibers.

The list of compounds identified as *kokumi* substances is growing (e.g. Amino et al, 2016). We tested 5 of these, including γEVG, glutathione (GSH), cinacalcet, γ-Glu-Abu, and CaCl_2_. All these compounds, applied orally, evoked responses in some trigeminal ganglion neurons. The response properties were mostly small (low amplitude), with variable latencies and prolonged durations. However, when tested sequentially (admittedly in a limited sample of neurons), the above 5 compounds did not stimulate a singular coherent population of trigeminal ganglion neurons. This suggests that if *koku* sensations are indeed transmitted via somatosensory afferent of the trigeminal ganglion, there is not a uniform group of “*koku*-responding” neurons in that ganglion. Moreover, the relatively small population of trigeminal ganglion neurons dedicated to oral stimulation with *kokumi* substances (for instance compared with thermal-orosensitive neurons, Leijon et al 2019) suggest that it will remain a challenge to study *koku*-evoked somatosensations at the cellular level in the trigeminal ganglion. In particular, relating *koku*-responsive neurons to molecular markers and eventually identifying a molecular receptor in trigeminal neurons will be challenging.

It is worth noting that *kokumi* substances also have been shown to activate CaSR-expressing taste bud cells (Maruyama et al, 2012). These cells are innervated by afferent fibers from the geniculate ganglion. Indeed, taste buds in the tongue and palate are associated with fibers from both geniculate and trigeminal ganglia (Fig. 13). Neurons from the geniculate ganglion innervate taste bud cells that are stimulated by taste substances entering through an apical taste pore. Thus, geniculate ganglion neurons typically are thought to transmit basic taste qualities such as sweet, umami, etc. In contrast, axons from trigeminal ganglia penetrate the oral epithelium immediately surrounding taste buds, forming a halo of nerves around most taste buds. Importantly, trigeminal fibers frequently are located in very superficial layers of the epithelium, where they may be reached by oral stimuli such as γEVG. In summary, trigeminal and geniculate ganglion neurons are positioned to respond to oral stimulation with *kokumi* substances – trigeminal neurons by direct stimulation, and geniculate neurons following stimulation of taste bud cells. The sensory characteristics of *kokumi* compounds are diverse, and it is thus possible that the trigeminal and geniculate ganglia play different, or complimentary, roles in the multifaceted taste that is *koku.*

**Figure 13.**
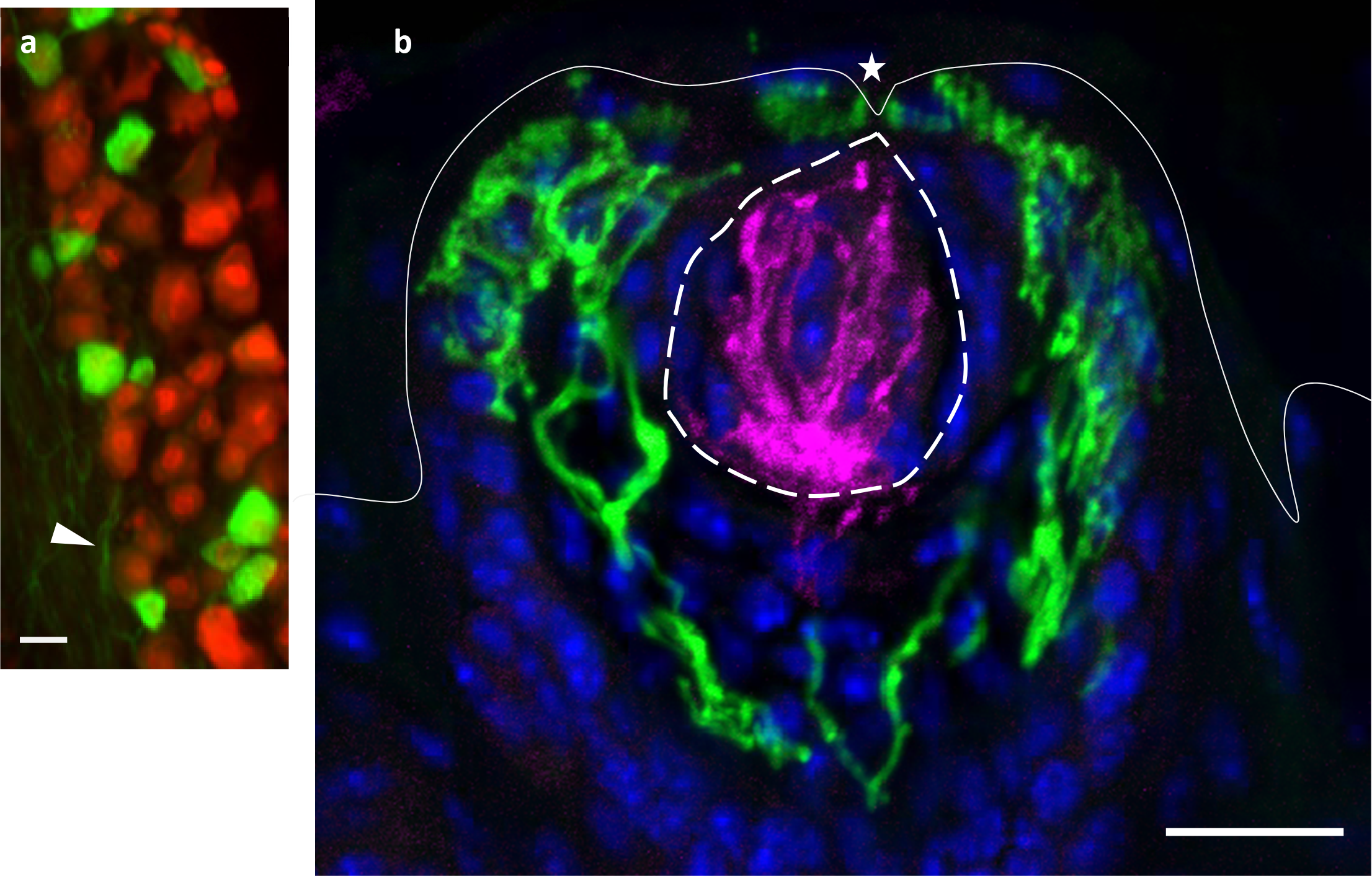
Afferent sensory fibers from trigeminal ganglion neurons terminate in the epithelium immediately surrounding taste buds. a) high magnification of a region of the trigeminal ganglion from a *Trpm8*-GFP mouse (Dhaka et al, 2007), immunostained for NeuN (red) to label neurons and GFP (green) to mark those neurons that express TrpM8. A subset of the neurons expresses GFP/TrpM8. GFP is also present in axons from these neurons (arrowhead). The geniculate ganglia from the same mouse, processed in parallel, did not contain GFP/TrpM8 (not shown). b) confocal micrograph of a fungiform papilla (solid outline) with a taste bud (dashed outline) in the tongue of the same *Trpm8*-GFP mouse. GFP/TrpM8-expressing axons surround, but do not enter the taste bud. In contrast, afferent fibers from the geniculate (gustatory) ganglion are immunoreactive for P2X2 (magenta) branch and branch extensively throughout the taste bud. The taste pore (*), through which taste stimuli interact with taste bud cells, is indicated (*). Scale bars, 20μm.

## Acknowledgements

This work was supported by grants from Ajinomoto Co., Inc., and the USA National Institutes of Health: NIDCD R01DC014420 (SR,NC), NIDCR/NCI R21DE027237 (SR). The authors thank Dr. Ajay Dhaka, University of Washington, for the generous donation of tissue used to produce Figure 13.

